# Second Climate Survey of Biomedical PhD Students in the Time of Covid

**DOI:** 10.1101/2022.01.13.476194

**Authors:** Deepti Ramadoss, John P. Horn

## Abstract

In July 2021, sixteen months into the Covid-19 pandemic, the institutional climate for PhD training in the School of Medicine was assessed for a second time. This survey of graduate students occurred 1 year after initial surveys of graduate students and training faculty in July 2020. The 2021 survey was completed by 99 PhD students in 11 PhD-granting programs. To allow comparisons between years, most of the 2021 questions were repeated with only minor edits. A few items were added to assess impacts of school-wide town hall meetings, a new PhD career club program, and enlarged mental health services. Several themes emerged. Students remain extremely concerned about the pandemic’s impact upon their training and long-term career prospects. They worry specifically about pandemic related reductions in research productivity and networking opportunities. Many students successfully adapted to laboratory research under pandemic restrictions but suffer from the continuing lack of social interaction even after in-person work hours increased. Symptoms of anxiety and/or depression persist amongst 46% of the students, as compared to 51% in 2020. Nearly 80% of students continue to report strong satisfaction with mentoring relationships with their dissertation advisors, but to lesser extents with programs (66%), departments or centers (71%), the School of Medicine (32%) and the University (49%). Students (26%) express interest in the Ombuds office that was announced in late 2021. Some students wrote that the medical school could do a better job in embracing diversity and inclusion and in mentor training, and many stated that town hall meetings do not serve them well. Coping mechanisms shared by some students demonstrate impressive resilience. These results present a mixed picture. While aspects of biomedical PhD training have begun to recover as the pandemic continues, long-term consequences of the disruption raise challenges that must be addressed by efforts to restore and improve the learning environment required for 21^st^ century research education.

## Background and Goals

The overarching goal for conducting a climate survey of biomedical PhD students in our school remains largely unchanged from that described in last year’s report [1]. Looking to the future, biomedical PhD training now requires mindful preparation for a diversity of possible career paths inside and outside of academia. Today’s biomedical PhD student must learn not only to conduct research using rigorous and reproducible approaches, but also to work effectively with colleagues having diverse personal and professional backgrounds, to build professional networks, to build personal wellness and professional resilience, and to engage in outreach to multiple types of audiences. The great majority of PhD students will follow a career path different from that of their dissertation mentor, a point documented by the landmark 2012 NIH workforce report [2] and from work through the NIH BEST program [3]. These trends are also evident in career outcomes of trainees in our school. Although some mentors do magnificent work in guiding their trainees along pathways to career success, many do not engage in efforts beyond the scientific goals of their specific research field. This approach was once widespread and considered adequate but no longer remains tenable. NIH and other funding agencies now require more comprehensive approaches to training and to the documentation of outcomes [4]. It therefore becomes imperative that research training programs and the institutions that house them establish resources to support career exploration and planning, wellness and resilience, diversity and inclusion, and mentor training. Together these elements can create a learning environment conducive to the success and satisfaction of all stakeholders. Assessing the institutional climate provides a means for understanding the learning environment. It can help to evaluate the success and effectiveness of strategies and interventions designed to improve learning.

When the Covid-19 pandemic exploded in March 2020, it disrupted all educational programs and research at the University of Pittsburgh together with our plans for a first climate survey of biomedical PhD students in the School of Medicine. Consequently, the 2020 survey was modified to include questions on how the pandemic was affecting students. In the 2021 survey, the goal of documenting climate during the pandemic continues to be of paramount importance. Although it may be too early to identify definitive trends, the data from this year and last provide baseline evidence for assessing future progress towards better training, career preparation and student satisfaction.

## Methods

The 2021 student survey contains 54 items, divided into groups labeled as Consent, Demographics, Wellness and Resilience, Mentoring, Career Goals, Career Exploration and Planning, Overall Satisfaction. Some items have multiple components and others provide open-ended comment spaces. The survey was constructed and implemented using Qualtrics^XM^ (Provo, UT), under the University of Pittsburgh license. The survey was approved by the University of Pittsburgh Institutional Review Board (IRB), and was sent on July 8, 2021 to all PhD students registered in School of Medicine programs, and to students in programs administered jointly with the Dietrich School of Arts and Sciences and with Carnegie Mellon University. Students received three reminders between July 16 and July 31.

Participation in the survey was voluntary. Participants gave informed consent before beginning the survey, which consisted of multiple-choice questions, and text boxes to make comments and/or elaborate on responses. Each participant was asked to provide their name, and the name of their dissertation advisor. The purpose of collecting this information is to permit longitudinal analysis of individual student trajectories at later time points in their careers, and to verify that each respondent was unique. After removing duplicate submissions, responses were anonymized using random alpha-numeric codes. In subsequent analysis the authors were blinded to student identity.

## Results and Discussion

The survey was completed by 99 of 367 PhD students, a return rate of 27%. Recipients included all 329 PhD students registered in the School of Medicine and 38 students enrolled in programs jointly administered with other schools. These included 31 students from the Neuroscience and the Molecular Biophysics & Structural Biology joint programs who were registered in the Dietrich School of Arts and Sciences, and 4 Molecular Biophysics & Structural Biology students who were registered at Carnegie Mellon University. Only 3 of the 38 students registered in other schools completed the survey. Thus 97% of the 99 respondents were registered in the School of Medicine.

### Demographics

Categories for self-identification of race/ethnicity were based on those used by NIH to track grantees and trainees. Of the survey respondents, 58% identified as White, 22% as Asian, 10% as Hispanic/LatinX, 7% as Black, and 2% as Native American, Hawaiian, Other Pacific Islander. Eight percent of these identified as more than one race, while 1% did not report race/ethnicity. These demographics resemble those for all 329 PhD students who were enrolled in the School of Medicine in July 2021: 50% White, 22% Asian, 8% Hispanic/LatinX, 5% Black/African American, 3% Multi-Racial, and 12% not specified.

As to gender identity, 56% of survey respondents identified as women, 40% identified as men, and 4% identified as gender queer, more than one identity, or did not respond. These statistics resemble the sex distribution of the entire school population of PhD students where 57% reported themselves as female https://somgrad.pitt.edu/graduate-data-dashboard.

Using their year of program entry, respondents were classified either as **early-stage PhD trainees** or **late-stage PhD trainees**. Early-stage trainees were defined as having completed one or two years of their program while late-stage trainees were defined as having completed three or more years of training. Note that July, when the survey was administered, is prior to matriculation of entering students in August – thus all respondents had at least one year of training and were familiar with topics raised by most survey questions. The data showed that 55% of respondents were early-stage trainees, while 46% were late-stage trainees. This differs from all SOM enrollees in July 2021 where 49% were early-stage trainees, and 51% were late-stage trainees. Program descriptions can be found at https://somgrad.pitt.edu/.

Students from all 11 PhD-granting programs responded to the 2021 survey (Table 1).

**Table 1.** PhD Program Affiliation.

| Table 1 – PhD Program Affiliation | % of survey respondents | Program as % of school's total enrollment |
| --- | --- | --- |
| Biomedical Informatics | 7 | 2 |
| Cell Biology & Molecular Physiology | 2 | 3 |
| Cellular & Molecular Pathology | 14 | 11 |
| Clinical & Translational Science | 1 | 2 |
| Computational Biology | 2 | 11 |
| Integrative Systems Biology | 6 | 10 |
| Microbiology & Immunology | 17 | 23 |
| Molecular Biophysics & Structural Biology | 2 | 3 |
| Molecular Genetics & Developmental Biology | 6 | 6 |
| Molecular Pharmacology | 17 | 12 |
| Neurobiology/Neuroscience | 14 | 17 |
| Blank | 1 |  |

The data in Table 1 indicate that variations in response rates generally reflect the size of programs. A similar trend was observed in 2020. Survey respondents also included 6 MD/PhD students who were enrolled as graduate students in the School of Medicine. This represents 18% of the 34 MD/PhD students enrolled as graduate students in the medical school PhD programs.

#### Written comments about the demographic questions

Some students asked about the need for demographic data. The reason is to provide a basis for determining whether different groups of students experience differences in the learning environment associated with gender, race, ethnicity, or stage of training. Other questions from respondents related to whether the categories of gender identity were adequate. In this case, the goal was to maximize inclusiveness while avoiding the creation of subgroups that were too small to include in the analysis and that might possibly compromise an individual’s identity in published reports.

Additional comments also asked about the survey’s lack of anonymity. As noted in the Methods, there are at least two reasons for this approach. The first is to detect and remove duplicate submissions. In the 2021 data, 3 such duplicates were found. The second more important reason is to permit longitudinal tracking of outcomes. Can one detect associations between responses to a climate survey and long-term career outcomes? Although difficult, this may prove important for assessing different components of the training environment and efforts to strengthen training. To protect the identity of individual respondents, all the data was anonymized and then analyzed blind (see Methods). This approach is analogous to clinical studies of identified subjects where personal information is kept confidential and the data is carefully protected. Identifying personal data are never disclosed in analyses, presentations or publications. The reliability of this approach depends upon the integrity of the investigators and adherence to the ethical standards overseen through the Institutional Review Board (IRB). The survey protocol for this work as well as the consent signed by respondents was reviewed and approved by the IRB at the University of Pittsburgh. The purpose of the climate survey was not to target individuals engaged in unacceptable mistreatment of students. If a student encounters such issues, there are three different routes to resolution:

1. A student may try to resolve the issue by talking with their advisor, program director or the Associate Dean for Graduate Studies.
2. If a student seeks confidential advice on a delicate problem, help is now available through the School of Medicine Ombuds office https://www.medschool.pitt.edu/ombuds-office and its Mental Health Team https://www.medstudentaffairs.pitt.edu/contact-us/mental-health-team.
3. Finally, if a student wishes to report instances of laudable or bad behavior, this can be done through the Office of the Learning Environment PAIRS system https://www.ole.pitt.edu/accolades/submit-accolade https://www.ole.pitt.edu/incidents/submit-incident-report.

### Wellness and Resilience

Graduate school and careers in biomedical science together with the uncertainties associated with young adult life can be inherently stressful [5]. Beyond these stresses and pressures, graduate students now must contend with the pandemic disruption of their studies and with extreme social isolation resulting from ‘distancing’, lock-downs and closures. Questions in this section were designed to probe work habits and to learn how students deal with the stresses they face. **The actual language of survey questions is italicized in blue**. Responses to matched questions from the 2020 survey are reported for comparison.

Although 99% of graduate students find time for recreational activities, 47% indicate it is insufficient (Table 2).

> *Do you find time for activities outside your studies and research? (examples include recreation, exercise, hobbies, social life, movies, music, theater, cooking)*

**Table 2.** Time for outside activities.

| **Table 2 – Time for outside activities** | % of responses 2021 | % of responses 2020 |
| --- | --- | --- |
| Yes | 52 | 58 |
| A little, but not enough | 47 | 37 |
| No | 1 | 5 |

In the 2020 survey, students were asked how much time they spent in the laboratory before the pandemic shut down in March 2020. In 2021 they were asked about their lab time after the pandemic shut down ended. The data indicate that students spent less time in the lab than before the pandemic (Table 3). Those spending 20 hours per week in lab increased from 5% before the pandemic to 25% in July 2021. Those spending 50 or more hours per week in lab dropped from 58% before the pandemic to 25% in July 2021.

> *During the past year, how many hours per week did you spend in the lab after the complete shut-down ended? Choose the answer that is closest to your schedule*.

**Table 3.**
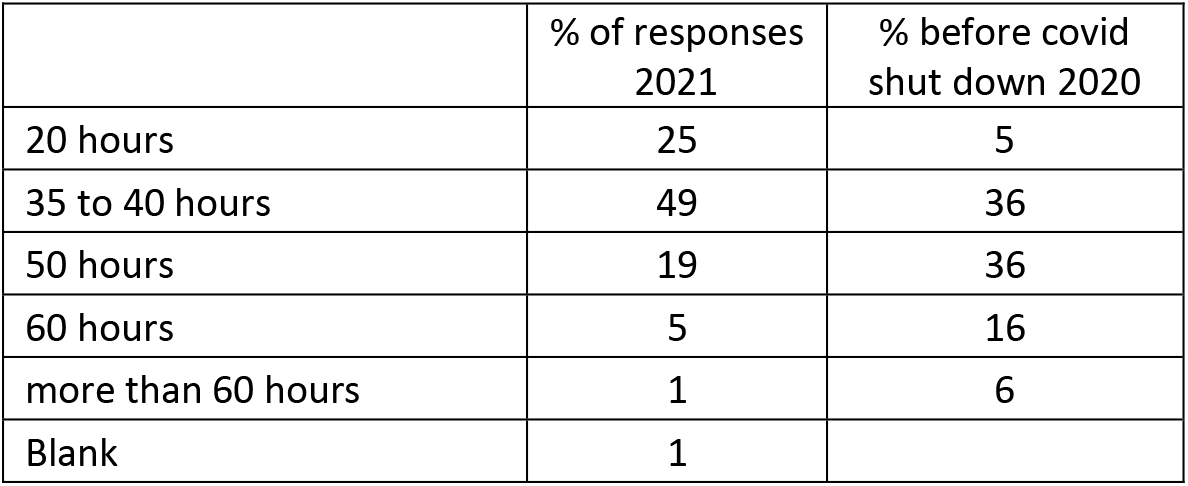
Lab hours/week.

The reduction in lab time seen in 2021 appears to be counterbalanced by changes in remote work since 2020 (Table 4). Students spending 20 hours/week in remote work increased from 28% to 59% while those working 35 to 45 hours/week in remote work decreased from 54% to 29% and those working 50 or more hours at home decreased from 18% to 12%.

> *During this past year, how many hours per week did you spend on average working remotely from home?*

**Table 4.** Remote hours/week.

| **Table 4 – Remote hours/week** | % of responses 2021 | % of responses 2020 |
| --- | --- | --- |
| 20 hours | 59 | 28 |
| 35 to 45 hours | 29 | 54 |
| 50 hours | 9 | 12 |
| 60 hours | 3 | 3 |
| more than 60 hours | 0 | 3 |

Working in the laboratory on weekends recovered to pre-pandemic behavior (Table 5).

> *During the past year, how often have you worked in the lab on weekends?*

**Table 5.** Working in the lab on weekends.

Working in the laboratory on weekends recovered to pre-pandemic behavior (Table 5). *During the past year, how often have you worked in the lab on weekends?*
| **Table 5 – Working in the lab on weekends** | % of responses 2021 | % before covid shut down 2020 |
| --- | --- | --- |
| Never | 11 | 8 |
| Sometimes | 48 | 42 |
| About half the time | 18 | 25 |
| Most of the time | 16 | 18 |
| Always | 7 | 7 |

The imposter syndrome can reflect a lack of self-confidence that often acts as a source of stress and anxiety in students. As seen in the 2020 survey, 50% of the students experience imposter syndrome half the time or more (Table 6).

> *Sometimes students say, “I feel like an impostor. Everyone else is smarter and I was admitted to the program by mistake.” Do you ever feel this way?*

**Table 6.** Feeling the Imposter syndrome.

| **Table 6 – Feeling the Imposter syndrome** | % of responses 2021 | % of responses 2020 |
| --- | --- | --- |
| Never | 21 | 13 |
| Sometimes | 29 | 37 |
| About half the time | 17 | 15 |
| Most of the time | 20 | 21 |
| Always | 13 | 14 |

The GAD-2 and PHQ-2 are pairs of questions in the survey that are used widely to screen for possible clinical anxiety and depression [6,7].

> *Over the **<u>last 2 weeks</u>**, how often have you been bothered by the following problems?*

Interpreting responses (Table 7) requires that each question be scored on a scale of 0-3 and then summed for each person for each pair of questions. This yields summed scores of 0-6 summarized in Table 8.

**Table 7.**
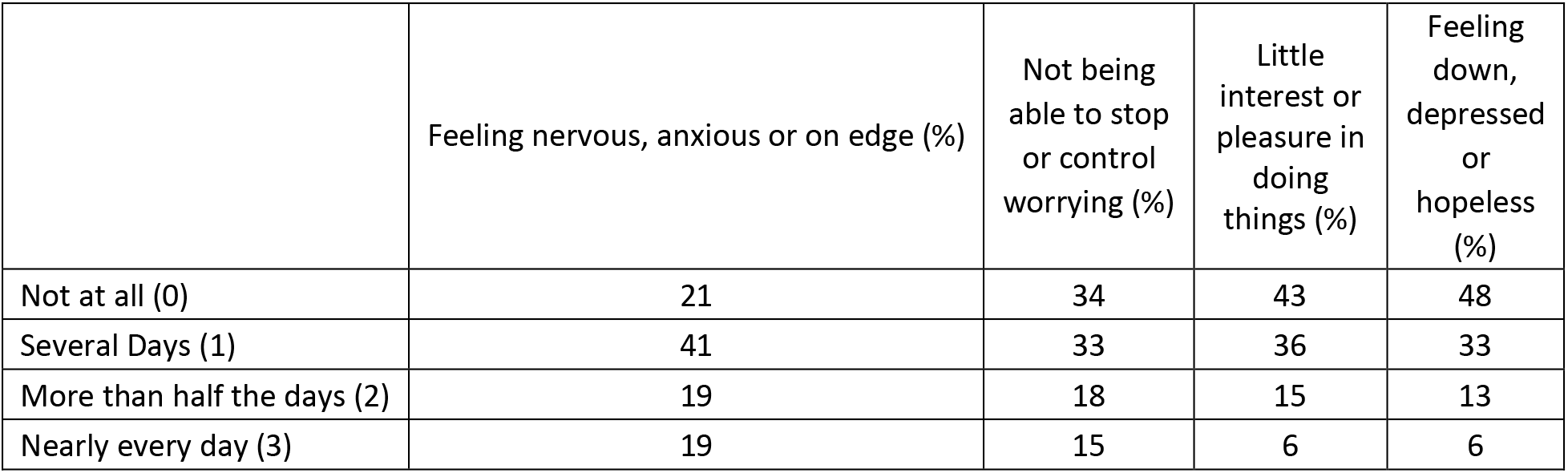
GAD-2 & PHQ-2 Questions and scores.

| **Table 7 – GAD-2 & PHQ-2 Questions and scores** | Feeling nervous, anxious or on edge (%) | Not being able to stop or control worrying (%) | Little interest or pleasure in doing things (%) | Feeling down, depressed or hopeless (%) |
| --- | --- | --- | --- | --- |
| Not at all (0) | 21 | 34 | 43 | 48 |
| Several Days (1) | 41 | 33 | 36 | 33 |
| More than half the days (2) | 19 | 18 | 15 | 13 |
| Nearly every day (3) | 19 | 15 | 6 | 6 |

**Table 8.** Total Scores.

| **Table 8 – Total Scores** | 0 | 1 | 2 | 3 | 4 | 5 | 6 |
| --- | --- | --- | --- | --- | --- | --- | --- |
| GAD-2 anxiety | 16 | 21 | 22 | 7 | 13 | 10 | 10 |
| PHQ-2 depression | 36 | 16 | 21 | 11 | 9 | 2 | 4 |

Table 8 shows the distribution of summed individual scores for each test (% of respondents).

A score ≥ 3 (shaded) on either test indicates a likelihood of clinically significant anxiety or depression that may require further evaluation. Table 9 shows the percentage of graduate students who had a score ≥ 3 on either test, or on both tests. Comparing the data for 2020 and 2021 reveals a small reduction in scores this year in all categories. The data also indicate that 46% of respondents scored ≥ 3 on either one or both tests in 2021, compared to 51% in 2020. While these results may signify a small improvement in the overall mental health of students, additional factors suggest otherwise. In July 2021, the vaccinations had come into effect, the pandemic toll of infections and death had declined and there was optimism about future improvement. Just as the survey was administered, the Covid Delta wave was beginning, now of course being followed by the Omicron wave. It is also clear from student comments (below) that the pandemic has exacted a high toll on student health that may not be adequately reflected by GAD/PHQ screening data. The long-term impact of the pandemic on the mental health of graduate students is concerning and cannot be overstated – in some cases, symptoms displayed by graduate students resemble those of PTSD [8].

**Table 9.** GAD/PHQ scores ≥ 3 (% of respondents)

| **Table 9 – GAD/PHQ scores ≥ 3**<br>**(% of respondents)** | **2021** | **2020** |
| --- | --- | --- |
| GAD-2 ≥ 3 (anxiety) | 40% | 46% |
| PHQ-2 ≥ 3 (depression) | 26% | 30% |
| Both GAD-2 and PHQ-2 C | 18% | 24% |

Despite efforts to strengthen and publicize the wellness resources available to all students at the University, only a small proportion of survey respondents utilize mental health services through the University counseling center – only 16% answered this question in 2021, which is unchanged since 2020 (Table 10).

> *Over the past few years, the University has invested in strengthening the counseling center. Have you ever used any of the services they offer? Check all that apply*.

**Table 10 -.**
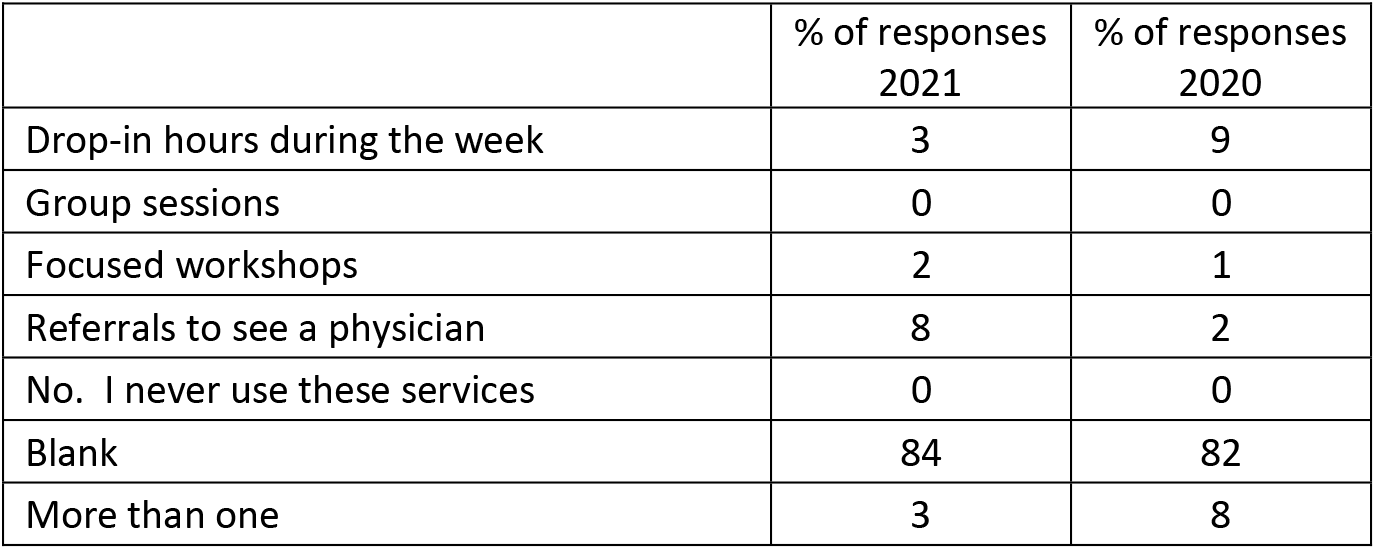
Counseling Center Utilization.

An Ombuds person can provide another mechanism to support the wellbeing of students. Prior to the pandemic, an Ombuds person for graduate students in the medical school was established and listed on the Provost’s website https://www.gradstudies.pitt.edu/about/school-ombudspersons. In late 2021, at Dean Shekhar’s direction a formal Ombuds office was created in the School of Medicine to serve the needs of all students. (https://www.medschool.pitt.edu/ombuds-office). When asked about their interest in an Ombuds office in the 2021 survey, 26% of graduate students indicated they were likely to use the Ombuds office. This was down from 37% in 2020, but still confirms the potential utility of this mechanism (Table 11).

> *The School of Medicine is in the process of creating an Ombuds office available to all students. Ombuds people are familiar with graduate education and the policies and procedures that govern them. The Ombuds Office is independent of the decision structure that evaluates students. Instead Ombuds people serve as a resource where students can go for confidential advice about their problems and concerns. How likely is it that you may want to speak with the SOM Ombudsperson?*

**Table 11.** Interest in an Ombuds Office.

| **Table 11 – Interest in an Ombuds Office** | **% of responses**<br>**2021** | **% of responses**<br>**2020** |
| --- | --- | --- |
| Extremely likely | 6 | 6 |
| Somewhat likely | 20 | 31 |
| Neither likely nor unlikely | 25 | 30 |
| Somewhat unlikely | 30 | 19 |
| Extremely unlikely | 16 | 14 |
| Blank | 2 |  |

Programs that embrace diversity and inclusion can exert a significant positive impact upon the overall culture and learning environment for graduate students in the School of Medicine. When asked whether such an embrace happens, respondents gave a mixed view, with small improvements between this year and last (Table 12). Positive support for diversity and inclusion was perceived by 58% of respondents in 2021, up from 53% in 2020. Nonetheless., 40% of respondents in 2021 continued to indicate that the graduate programs show little or no concern for diversity and inclusion, albeit down from 47% in 2020.

> *To what extent do graduate programs in the School of Medicine have a culture that embraces diversity and inclusion?*

**Table 12.** Embrace of Diversity & Inclusion by graduate programs.

| **Table 12 – Embrace of Diversity & Inclusion by graduate programs** | % of responses<br>2021 | % of responses<br>2020 |
| --- | --- | --- |
| not at all | 10 | 10 |
| a little | 30 | 37 |
| sometimes | 35 | 35 |
| definitely yes | 23 | 18 |
| Blank | 2 |  |

PhD students play an essential role in advancing the research mission of the medical school, a mission that is highly valued for its impact on the school’s reputation. It thus becomes important to understand whether graduate students feel recognized for their contribution to the success of the academic community (Table 13). While 69% of respondents in 2021 indicate they feel welcome as members of the community, 31% do not. The numbers did not change since 2020.

> *People like me feel like they are welcome as members of the larger School of Medicine community*

**Table 13 -.** Welcoming community.

| **Table 13 - Welcoming community** | % of responses<br>2021 | % of responses<br>2020 |
| --- | --- | --- |
| not at all | 8 | 5 |
| a little | 23 | 25 |
| sometimes | 28 | 31 |
| definitely yes | 39 | 39 |
| Blank | 2 |  |

Beyond education, the pandemic brought to light many glaring problems in American society that emanate from inequity and racism. In response to this awakening, graduate students and medical students began questioning what the School of Medicine could do to continue their training while also advancing social justice. To address these issues, while also improving communication and giving students a louder voice, Vice-Dean Thompson initiated a series of **monthly town hall meetings with students**. The meetings are attended by all deans involved in education within the school. This approach is highly unusual and remains so. Most medical schools do not hold meetings that bring together graduate students and medical students. The 2021 climate survey therefore asked for written feedback on the town halls.

> *Since June 2020, Vice-Dean Thompson has hosted monthly town hall meetings for all students in the School of Medicine. Have you attended any of these meetings? What have you found useful? What could be better?*

Thirty students responded that they attended one or more of the town halls. The most common view was that the town halls focused on medical student issues with very little discussion for or by graduate students. Several respondents noted that the meetings were overly long, not well-structured, and frustrating because they often did not articulate clear solutions. On the positive side, it was pointed out that having breakouts for medical and graduate students was helpful. Several students said they appreciated information about the pandemic discussed at town halls. A couple of students asked why only deans, but not other faculty members were present. Although it is true that some faculty attended the first town halls, later meetings were restricted to Deans. The reason was to provide more space for students to express their views. Even so, some respondents expressed frustration that Deans talked too much and did not give enough time to hear student concerns.

One of us (JPH) has attended the town halls and appreciates all the student feedback. The meetings have taken an important first step in providing a forum for all students in the School of Medicine to be together to address common issues. We must recognize that although breaking out into silos may be useful for addressing some problems, it is not the total solution. It is difficult, yet important that we find areas of common ground that affect us all while also recognizing distinctions with areas more specific to scientific and medical training. While these meetings may be flawed, we also must each recognize that none of us has lived through challenges like those presented by the pandemic, making it a messy problem to work through.

All of us – scientist, physician or other – harbor bias. Bias arises at the subconscious level; it is a by-product of an important brain function to make decisions with incomplete information (examples include optical and auditory illusions) to aid human survival. Bias can shade our conscious interpretation of experimental and clinical data; it can cause us to use limited knowledge to make illusory judgements about other people as individuals or groups, feeding racism and inequity. Although bias has a rational explanation, it does not have a rational justification. We as scientists and physicians must work to limit the influence of bias, by becoming more self-aware and working to mitigate the negative impacts of bias in our professional and personal lives. For this reason, the biomedical community has come to recognize the fundamental importance of bias training, especially the unconscious or implicit kind. To achieve this goal, constituencies across the university have worked to promote bias training. We therefore asked about it in the climate survey (Table 14).

> *How many times have you participated in a workshop or online training on the topic of implicit bias?*

**Table 14.**
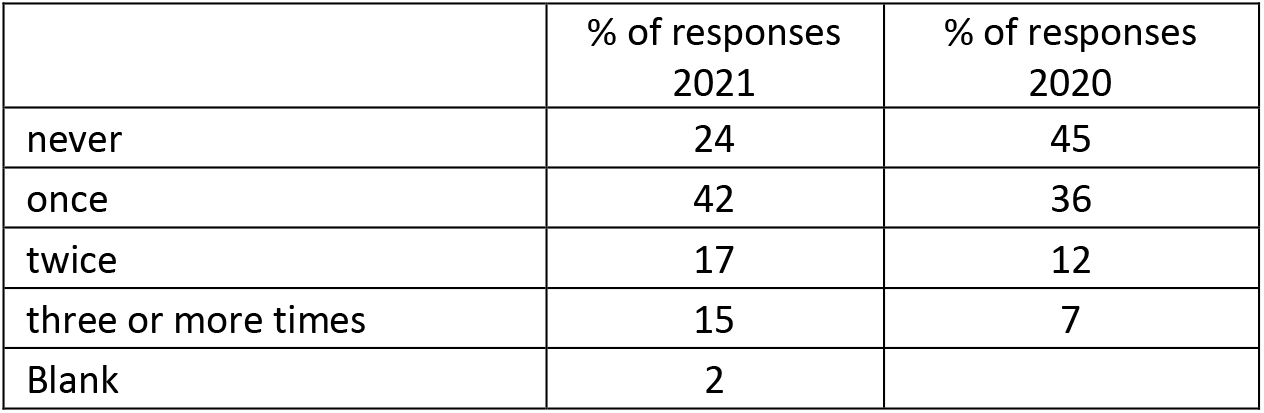
Frequency of implicit bias training.

Every aspect of bias training showed improvement over the past year. Respondents who had never been trained dropped from 45% in 2020 to 24% in 2021. All other categories from one training experience to three or more saw increases. The goal is for all trainees to be trained about bias, and the survey results suggest that the graduate programs are on the right track.

Bias and racism are often expressed through microaggressions. Although sometimes subtle such behaviors exert a pernicious impact that is incompatible with the climate required for a productive and just learning environment. When asked about microaggressions the responses remain difficult to interpret (Table 15). The incidence of individuals who never receive or witness microaggressions was 29% in 2021, down from 39% in 2020. However, the overall incidence of such behaviors increased to 69% in 2021, from 61% in 2020. It remains unclear whether these data reflect an increase in incidence of microaggressions or greater awareness of when such incidents happen. Resolving this question will require further study.

> *How often have you been exposed to microaggressions in the context of your graduate education at Pitt? - Either on the receiving end or as a witness?*

**Table 15.** Recipient or witness to microaggressions.

| **Table 15 – Recipient or witness to microaggressions** | **% of responses**<br>**2021** | **% of responses**<br>**2020** |
| --- | --- | --- |
| never | 29 | 39 |
| once | 8 | 5 |
| a couple of times | 41 | 32 |
| more than 5 times | 16 | 21 |
| all the time | 4 | 3 |
| Blank | 2 |  |

The survey then asked for-

> *Written comments concerning issues of wellness, resilience, diversity, and inclusion? **We are especially interested in how the pandemic impacted all aspects of your wellness and resilience.***

The responses were voluminous and ranged widely from very positive to very painful. Some respondents indicated they were doing very well and thankful for support provided through the medical school and its programs. Others focused on the painful effects of the pandemic and accompanying isolation they have endured through separation from friends and loved ones. Comments centered around multiple themes summarized below, with edited examples.

- The lack of racial and gender diversity, including attitudes regarding varied political viewpoints, highlight the need to address these issues in the community to improve climate. Examples:
  - The SOM could do better in promoting diversity.
  - We need more women and people of color in leadership positions.
  - The environment is intolerant of ideological diversity
- The pandemic added stress over progress and affected mental health in many ways. Some students commented that mental health support was inadequate. Examples:
  - My advisor has no concern about how the pandemic affected me, my family, my mental health or my work
  - Mental health access is a joke.
  - Individual programs could have done more to promote mental health resources
  - The University counseling center helped me learn ways to cope with my mental health issues.
- Several students described the loneliness caused by the pandemic, lack of personal interaction and the need for special support for first years and international students. Examples:
  - I felt incredibly isolated and alone, never knowing what was really going on.
  - Starting graduate school in a new city where I didn’t know anyone was tough.
  - The pandemic has been especially hard for international students, who have not been able to see their families, or are limited by travel and visa restrictions.
- Comments surrounding microaggressions, lack of support from faculty, programs, and higher administration were also prominent. Examples:
  - All microaggressions came from professors through comments about student performance and participation in zoom classes and sharing grades with the whole class.
  - Rare emails or support (if at all) from the medical school, the graduate studies office, department, or program are insufficient.
- Some students reported added expectations in the lab during the pandemic. This contrasted with a minority who found the ability to improve work-life balance with more opportunities for work from home. Examples:
  - The pandemic has morphed the expectations of work/research and has therefore put more stress and pressure on me.
  - I constantly feel on edge regarding expectations of “making up time” lost to the pandemic, imposed by my advisor and research group.
  - During the pandemic, it was harder to take breaks from lab because there wasn’t much else to do. I am trying to re-learn how to have a work-life balance.
  - Remote work was terrific for me and I hope it continues. I was more productive and this definitely helped with wellness because I found time to do other things.

Reading these comments, one cannot escape the conclusion that graduate students are paying a high price for the pandemic disruption and that the toll continued to rise during the second year of Covid. Feedback from the comments also suggest an action agenda of important issues to discuss with students in future town halls.

### Mentoring

Effective relationships between mentors and their mentees are seen as very strong drivers of academic success, research productivity, wellness, and satisfaction. Questions in this section probed mentoring relationships between graduate students and the training faculty.

Students are encouraged to develop secondary mentoring relationships with faculty members other than their thesis advisor. In the 2021 survey, 61% of PhD students reported having at least one secondary mentor (Table 16)

> *In addition to their dissertation adviser, students sometimes develop informal mentoring relationships with other faculty. Sometimes these secondary mentors are from one’s thesis committee. Other times they are from the student’s program, a journal club or are lab neighbors. How many secondary mentors do you have besides your dissertation adviser?*

**Table 16.** Number of secondary mentors.

| **Table 16 – Number of secondary mentors** | **% of responses 2021** | **% of responses 2020** |
| --- | --- | --- |
| 0 | 38 | 28 |
| 1 | 35 | 27 |
| 2 | 15 | 29 |
| 3 | 5 | 8 |
| 4 or more | 6 | 8 |

**Table 17.**
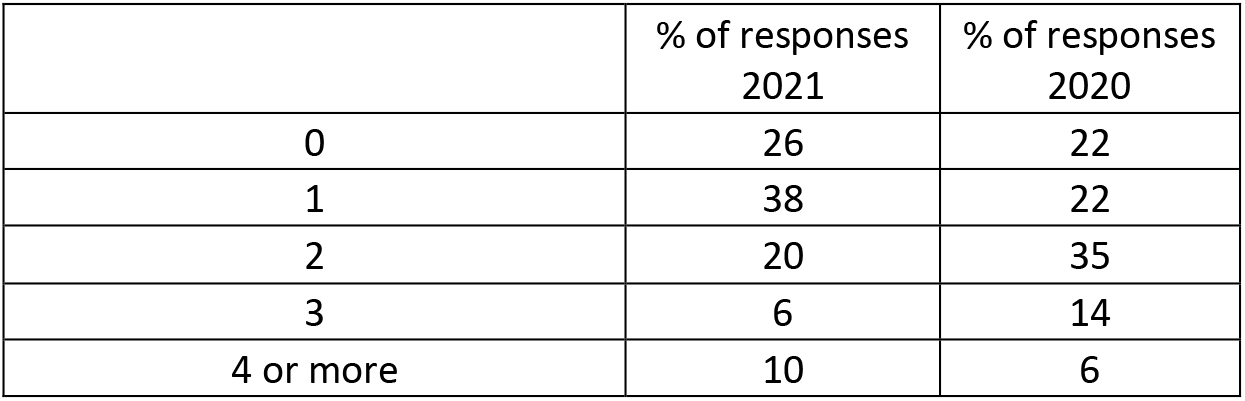
Number of secondary mentors for late-stage trainees.

The reduction in secondary mentors in 2021 from 2020 (i.e. of 10%) may have arisen from the 6% increase in the proportion of early-stage trainees in survey responses. One might imagine that mentoring relationships expand after students advance to candidacy. To explore this possibility, we compared secondary mentoring rates for late-stage trainees.

After removing the early-stage students, 74% of students report having at least one secondary mentor, while in 2020, 77% reported having more than one secondary mentor.

Effective PhD training requires regular one-on-one meetings between students and their advisors. The incidence of such meetings on a regular weekly or daily basis increased to 69% in 2021 from 58% in 2020 (Table 18). This paralleled a drop in less frequent meetings to 28% in 2021 from 38% in 2020. Although positive, the data suggest that programs must place more emphasis on frequent meetings between faculty members and their students.

> *On average, my dissertation adviser meets **individually** with me for 15 minutes or more*.

**Table 18.** Frequency of student meetings with their thesis mentor.

| **Table 18 – Frequency of student meetings with their thesis mentor** | **% of responses 2021** | **% of responses 2020** |
| --- | --- | --- |
| Almost every day | 7 | 4 |
| At least once a week | 62 | 54 |
| Every other week | 18 | 19 |
| Once a month | 7 | 13 |
| Less than once a month | 3 | 6 |
| Blank | 3 | 4 |

The AAMC compact [9] between graduate students and mentors is distributed to students during orientation and in a course on the Responsible Conduct of Research at the end of year one. The survey results indicate this approach is ineffective. Nearly 90% of respondents (Table 19) never use the compact, a figure that was unchanged between 2021 and 2020. This indicates that another approach will be required if the programs see the compact as a valuable resource.

> *The AAMC (American Association of Medical Colleges) publishes a compact to guide discussions between graduate students and their mentors. How many times have you used this compact in discussions with your dissertation adviser?*

**Table 19.**
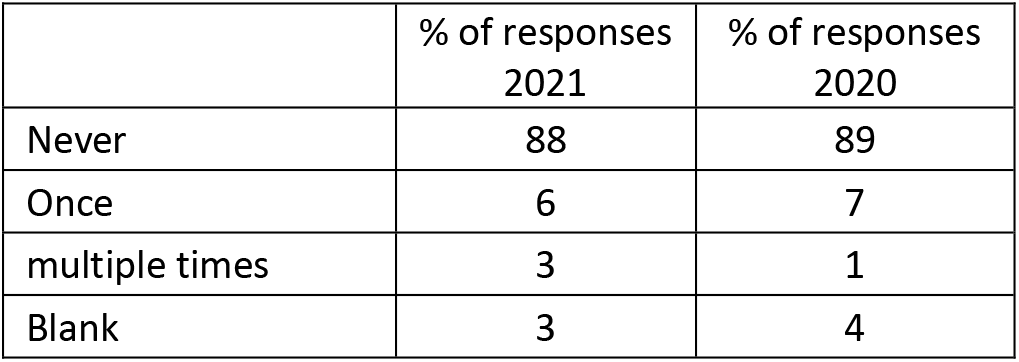
Use of the AAMC Compact.

Aside from the AAMC compact, the AAMC recently published the Appropriate Treatment of Research Trainees (AToRT) [10]. Use of this document for training sessions for mentors, new trainees, and departments are among its potential applications.

To probe perceptions of primary mentors, students were asked to rate them on a 0 −10 scale (weak to strong) in several areas describing their relationships.

> *My dissertation adviser cares about me as a person*
>
> *My dissertation adviser treats my scientific ideas and insights with respect*
>
> *My adviser gives me freedom to chart the direction of my dissertation research*
>
> *My dissertation adviser and I are on the same page when it comes to understanding each other’s mutual expectations*
>
> *When I have a problem, either personal or professional, my dissertation adviser displays empathy. Even when they cannot solve the problem, they are willing to listen*

Table 20 shows the full-scale data. To simplify comparisons, strong positive ratings were compiled by summing the percentage of responses giving a score of 7-10 (shaded) for each category. Using this approach, 77% of students feel their mentor cares about their personal well-being, 79% feel their scientific ideas are respected, 70% feel they have freedom to chart their own research direction, 68% feel their mutual expectations are matched, while 73% agree that their mentors display empathy. When compared to the 2020 data, the 2021 ratings are down by 2 to 10%. The reason for this change remains open to interpretation. Of greater concern is that 5 to 10% of respondents give their mentors a negative rating of 0 to 3.

**Table 20.** 2021 student ratings of mentors.

| **Table 20 – 2021 student ratings of mentors** | Cares about me as a person | Respects my scientific ideas | Freedom to chart research direction | Same page on mutual expectations | Displays empathy |
| --- | --- | --- | --- | --- | --- |
| 0 | 1 | 0 | 1 | 0 | 1 |
| 1 | 0 | 1 | 4 | 3 | 3 |
| 2 | 3 | 2 | 1 | 2 | 1 |
| 3 | 1 | 4 | 1 | 5 | 1 |
| 4 | 6 | 3 | 1 | 3 | 3 |
| 5 | 3 | 6 | 9 | 6 | 7 |
| 6 | 6 | 2 | 7 | 8 | 4 |
| 7 | 8 | 13 | 11 | 15 | 13 |
| 8 | 18 | 12 | 13 | 17 | 8 |
| 9 | 11 | 10 | 19 | 11 | 10 |
| 10 | 40 | 44 | 27 | 25 | 42 |
| Blank | 2 | 2 | 5 | 4 | 6 |
| Sum of 7-10 | 77 | 79 | 70 | 68 | 73 |
| 2020 Sum of 7-10 | 88 | 84 | 73 | 70 | 78 |

Verbal expressions of anger towards others in a professional academic environment constitute unacceptable behavior [10]. When asked about this, 82% of 2021 respondents (Table 21) reported that their mentors either do not engage in such behavior at all or did so only once. It remains a matter of serious concern that 14% of advisors behave this way on a repeated, sometimes weekly basis, and that this number that did not improve since the 2020 survey.

> *My dissertation adviser gets angry, raises their voice and yells at people*

**Table 21.** Expression of anger by mentors.

| **Table 21 – Expression of anger by mentors** | % of responses 2021 | % of responses 2020 |
| --- | --- | --- |
| never | 73 | 77 |
| once | 9 | 10 |
| occasionally | 12 | 12 |
| almost every week | 2 | 2 |
| Blank | 4 |  |

Mentors play a critical role in the professional development of their graduate students, this often happens by having them attend scientific meetings. It was a matter of concern that a large and increased proportion of students reported never presenting at a scientific meeting rising to 46% in 2021 from 28% in 2020 (Table 22).

> *How many national scientific meetings (virtual or in person) have you attended and presented an abstract? (since starting graduate school at Pitt)*

**Table 22.** Conferences attended per year.

| **Table 22 – Conferences attended per year** | % of responses 2021 | % of responses 2020 |
| --- | --- | --- |
| None | 46 | 28 |
| 1 | 15 | 24 |
| 2 | 12 | 23 |
| 3 | 12 | 8 |
| 4 or more | 12 | 17 |
| Blank | 3 |  |

Possible explanations for these data are the disruption of conferences by the pandemic, and a disproportionate impact upon early-stage trainees. We therefore examined conference participation by late-stage trainees who are more likely to have data suitable for presentation (Table 23). Although this accounts for some of the data, it remains a concern that 18% of late-stage trainees rarely attend conferences.

**Table 23.** Conferences attended by late-stage trainees per year.

| **Table 23 – Conferences attended by late-stage trainees per year** | % of responses 2021 | % of responses 2020 |
| --- | --- | --- |
| None | 18 | 8 |
| 1 | 16 | 25 |
| 2 | 18 | 32 |
| 3 | 20 | 9 |
| 4 or more | 27 | 24 |
| Blank | 2 | 3 |

Beyond simply allowing students to attend and present at meetings, mentors can actively encourage such participation and facilitate student participation in conferences. When asked about this (Table 24), 21% of respondents indicated that their mentors have <u>never</u> encouraged them at scientific meetings, up from 16% in 2020.

> *My dissertation adviser has encouraged and facilitated my participation in national scientific meetings*

**Table 24.** Mentor encouragement of student participation in conferences.

| **Table 24 – Mentor encouragement of student participation in conferences** | % of responses<br>2021 | % of responses<br>2020 |
| --- | --- | --- |
| Never | 21 | 16 |
| Occasionally | 33 | 29 |
| At least once a year | 42 | 55 |
| Blank | 3 |  |

Dividing the 2021 data into early-stage and late-stage trainees (Table 25) shows that most students who have never been encouraged to attend a meeting are in the first and second years. Nevertheless, the data does not explain why 4% of late-stage trainees are never encouraged and 14% are rarely encouraged to participate in meetings.

**Table 25.**
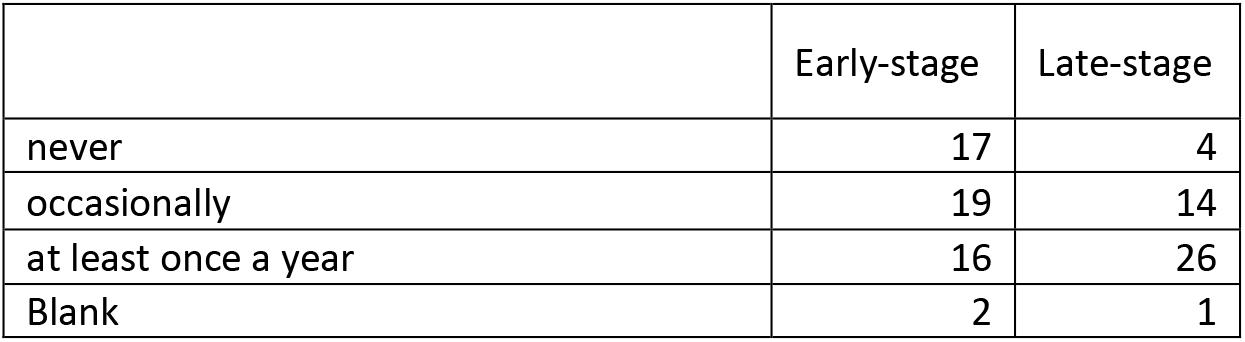
2021 Mentor encouragement of student participation in conferences.

When a student attends a conference with their mentor, it provides an important opportunity for the mentor to introduce the student to well-known established scientists in their field. This interaction can play a vital role in helping trainees to gain self-confidence and build their professional networks. When asked (Table 26) 43% of 2021 respondents report they have never been introduced at meetings to eminent people in their field, up slightly from 40% in 2020.

> *At scientific meetings, my dissertation adviser has introduced me to established scientists in my field who work at other universities*.

**Table 26.** Introductions at conferences to established scientists by mentors.

| **Table 26 – Introductions at conferences to established scientists by mentors** | % of responses<br>2021 | % of responses<br>2020 |
| --- | --- | --- |
| never | 43 | 40 |
| occasionally | 34 | 29 |
| at every meeting | 14 | 31 |
| blank | 8 |  |

Dividing the 2021 data into early-stage and late-stage trainees (Table 27) accounts for some instances where students are not introduced to other scientists. Nonetheless, 11% of late-stage graduate students have never been introduced to prominent scientists by their mentors, while 20% have only occasionally had this opportunity.

**Table 27.** 2021 Introductions to established scientists at conferences by stage.

| **Table 27 – 2021 Introductions to established scientists at conferences by stage** | Early-stage | Late-stage |
| --- | --- | --- |
| never | 32 | 11 |
| occasionally | 14 | 20 |
| at least once a year | 1 | 13 |
| blank | 7 | 1 |

Mentors play a critical role in training students to communicate their scientific ideas, especially through writing, which is essential for publishing research and receiving recognition. The survey therefore asked students to evaluate the help mentors have provided with scientific writing using an 11-point scale (Table 28).

> *My dissertation adviser has spent time helping me become a better scientific writer (totally true = 10, totally false = 0)*

**Table 28.** mentor support for writing skills & program requirements.

| **Table 28 – mentor support for writing skills & program requirements** | **Writing**<br>**% of responses**<br>**2021** | **Writing**<br>**% of responses**<br>**2020** | **Program requirements**<br>**% of responses**<br>**2021** | **Program requirements**<br>**% of responses**<br>**2021** |
| --- | --- | --- | --- | --- |
| 0 | 2 | 1 | 1 | 0 |
| 1 | 2 | 1 | 2 | 3 |
| 2 | 2 | 2 | 1 | 3 |
| 3 | 9 | 1 | 4 | 4 |
| 4 | 4 | 0 | 3 | 3 |
| 5 | 6 | 10 | 16 | 6 |
| 6 | 11 | 10 | 5 | 8 |
| 7 | 11 | 17 | 12 | 11 |
| 8 | 13 | 19 | 18 | 17 |
| 9 | 12 | 8 | 9 | 13 |
| 10 | 21 | 33 | 22 | 31 |
| Blank | 6 |  | 6 |  |
| Sum of 7 - 10 | 57 | 77 | 61 | 72 |

As in Table 20, ratings of 7 to 10 in Table 28 were considered as highly positive (green) and summed. Using this criterion, 57% of mentors in 2021 provided strong support with writing skills down from 77% in 2020. Negative ratings (0 – 3) were reported by 15% of respondents in 2021, up from 5% in 2020. Although the reason for this troubling change is unclear, it is too large to be accounted for by the shift in survey participation by early and late-stage trainees.

Another way that mentors promote the success of their trainees is by keeping them on track. Program structures provide a wholistic framework that goes beyond the dissertation project. The survey therefore asked students to rate their advisor’s knowledge of program requirements (Table 28). The respondents gave excellent ratings to 61% of mentors in 2021, down from 72% in 2020. Negative ratings (0-3) were given to 8% of mentors in 2021, down from 10% in 2020.

> *My dissertation adviser is familiar with the guidelines and milestones of my program and helps me stay on track. (totally true = 10, totally false = 0)*

### Career Goals

Students were asked to prioritize their long-term career goals with questions based on a condensed version of the three-tier taxonomy of career outcomes that was developed by members of the BEST consortium [8] and adopted by the Coalition for Next Generation Life Sciences (https://nglscoalition.org). The tiers describe “Workforce/Employment Sector”, “Career/Job Type”, and “Job Function”. Our questionnaire used one tier that contained 5 employment sectors (n=89 respondents) and a second tier with 11 job functions (n= 94 respondents).

Preferred employment sector based on student rankings were – 1) academia (33%), 2) business/for-profit (32%), 3) government (10%), 4) undecided (10%), and 5) other non-profit (4%). Table 29 shows the same order of preference in the 2020 survey data.

**Table 29.** Preferred employment sector.

| **Table 29 – Preferred employment sector** | % of responses<br>2021 | % of responses<br>2020 |
| --- | --- | --- |
| Academia | 33 | 36 |
| Business (for-profit) | 32 | 35 |
| Government | 10 | 5 |
| Non-profit | 4 | 3 |
| Undecided | 10 | 22 |

Students were also asked to prioritize their long-term job goals in rank order. Table 30 shows the top four choices in 2021 were 1) Research & Development in industry (31%), 2) Tenure track professor at an R1 university, research institute or government lab (25%), 3) Staff scientist in a research-intensive non-profit organization (11%), and 4) professor who teaches and has a lab at a small college (8%). This ranking differs from 2020 when the top choice was Tenure Track Professor (31%), and the second choice was R & D in industry (26%). Choices three and four remained unchanged - Staff Scientist (6%), and Professor with lab at a small college (5%).

**Table 30.** Preferred long-term job goal.

| **Table 30 – Preferred long-term job goal** | % of responses<br>2021 | % of responses<br>2020 |
| --- | --- | --- |
| Tenure-track professor in research intensive University, Research Institute or Government lab | 25 | 31 |
| Staff Scientist in research intensive organization | 11 | 6 |
| Teaching professor in a small college | 3 | 2 |
| Professor who teaches and has a lab in an undergraduate college setting | 8 | 5 |
| Research & Development in industry | 31 | 26 |
| Intellectual property, law, business of science | 3 | 4 |
| Scientific publishing | 1 | 3 |
| Science policy | 1 | 1 |
| Science sales + customer ed | 0 | 1 |
| Other | 4 | 8 |
| Undecided | 7 | 14 |

Students were also asked about their next step after completing the PhD (Table 31).

> *What is the **first position you will seek** immediately after graduate school? (OK to check more than one choice)*

As found in the 2020 survey, the first choice of most students in the 2021 survey remains postdoctoral training and the second most popular choice remains an entry level job in big pharma or biotech.

**Table 31.** Goal for first job after PhD.

| **Table 31 – Goal for first job after PhD** | **% of responses**<br>**2021** | **% of responses**<br>**2020** |
| --- | --- | --- |
| Postdoctoral research training | 47 | 42 |
| Entry level job in big pharma or a biotech company | 31 | 27 |
| Entry level job in publishing | 2 | 3 |
| Teaching job | 4 | 5 |
| Policy internship in government or a non-profit organization | 8 | 6 |
| Business or law school | 1 | 2 |
| Undecided at this time | 8 | 12 |
| Blank |  | 2 |

Several themes emerged from comments students shared on the effect of the pandemic on their job goals, long-term and short-term.

> *How were your goals impacted by the pandemic? What activities a helped you cope with pandemic-related disruptions?*

The most frequent comments focused on pandemic disruption of networking – particularly in the context of career advancement, as well as with peers. Many noted the negative impact of pandemic-related delays in research progress and graduation; this impact has also been observed at other institutions and in other academic fields [8]. Several early-stage students did not feel that their goals were immediately impacted due to the pandemic. Other students said their career goals had changed or matured, prompting them to consider moving away from academia. The reasons for the shift included stable jobs not dependent on grant funding, flexibility of virtual work, and attitudes expressed by professors during the pandemic. Issues not related to the pandemic were also shared, such as contending with issues in academia like conflicts of interest, rigor and reproducibility, and looking for employment outside of biomedical paths altogether. On the positive side, some students said the pandemic enhanced their ability to participate in internship opportunities, and to publish research.

In this section the pandemic toll on mental health is once again palpable. Some students describe their inability to think about their careers during the pandemic, and the impact on their mental health, while others shared coping with strategies such as exercise, reading, cooking, baking, non-profit work, online socials, and counseling.

### Career Exploration and Planning

When asked about career exploration and planning, 49% of students think about or plan for their futures all the time, or a good deal of the time, while 31% spend a moderate amount on career planning. The remaining 17% spend only minimal time on this type of planning.

Students were also asked about barriers to career exploration and planning that they believe they face, and were provided multiple options, and asked to select as many as applied to them – several students selected more than one barrier. The most common response selected by 34% of students was the idea that the task feels too overwhelming, with too many unknowns and possibilities. Coming in a close second, 33% of students cited the lack of time to spend on career planning. Only 5% of students perceived their advisor as not supportive of their interest in a career outside of academia. On a further positive note, only 4% of students reported that their advisors were pressuring them to spend more time in the lab, rather than be diverted by career planning. Also encouraging, 22% of students indicated they felt good about their plan, and that their advisors were supportive of their efforts.

From our career development interactions with students, we know that there is a fair amount of interest in internship or externship opportunities to explore career options. When asked specifically about this, 70% of students indicated moderate to extreme interest in such options, while 3% indicate they have already begun participating in these opportunities. The remaining 24% indicated they were either not interested or only slightly interested.

The School of Medicine Office of Graduate Studies promotes career and professional development opportunities organized by the office, and by partners, such as the Biomedical Graduate Student Association (BGSA), the Office of Philanthropic and Alumni Engagement (PAE), and the Office of Academic Career Development (OACD). Only 22% of students have participated in these activities frequently, while 24% have attended only one event. The remaining 51% have never taken part in these events, which is not surprising because 32% of this group are early-stage graduate students who are more focused on getting their PhD training off the ground.

When asked to name events they have attended, the most popular choices included career panels with alumni, panels of people working in different sectors, career sessions on specific topics, and career fairs. A couple of respondents also noted they had participated in the Career Club organized by the Office of Graduate Studies. The club is a longitudinal series of 6 workshops designed to include career explorations, networking, resume and interview preparation, and outreach to potential employers.

To gauge student interest in the Career Club, students were queried on their interest levels. 42% of the students responded they would be interested in participating when they are eligible (4^th^ year and beyond), while 14% were not interested and 5% had already participated. Those expressing possible interest (35%) stated that it would depend on their availability and time, research progress, topics covered, learning more about the program, and access to participate.

Students were also asked to comment on career planning and exploration. Students reported that they had heard positive things about the Career Club, were keen to participate. While some students felt they were on track in planning for the future, others stated it was too early to spend time on this topic. Some responses voiced concern over career readiness, how to find jobs, and credentials needed for jobs outside academia (e.g. are postdocs necessary?). In general, a majority of students are very interested in career development opportunities and concerned about the difficulty of navigating the process during the pandemic disruption. Their feedback suggests that efforts by the school to support these activities are on the right track.

### Overall Satisfaction

The final section of the survey asked students to rate their satisfaction with the different elements of their educational experience.

> *Putting it all together, I am happy with these elements of my training*

The data expressed as the percent of respondents giving each response (Table 32) indicate that clear majorities of students report being moderately or extremely happy with their PhD program (66%), dissertation advisor (78%), dissertation project (63%), and academic department or center (61%). By contrast, only 32% are extremely or moderately happy about being valued members of the SOM community, and only 49% are extremely or moderately happy about feeling a part of the larger university community. This mirrors a larger study where students reported feeling more support from their advisors than from their school or university [8].

**Table 32.** Overall ratings of student satisfaction (%)

| **Table 32 –**<br>**Overall ratings**<br>**of student**<br>**satisfaction**<br>**(%)** | PhD<br>Program | Dissertation<br>advisor | Dissertation<br>project | Department<br>or center | Valued member<br>of School of<br>Medicine<br>community | Part of the<br>University of<br>Pittsburgh<br>community |
| --- | --- | --- | --- | --- | --- | --- |
| Extremely<br>happy | 22 | 49 | 30 | 24 | 9 | 20 |
| Moderately<br>happy | 44 | 29 | 33 | 37 | 23 | 29 |
| Slightly happy | 15 | 3 | 16 | 15 | 21 | 19 |
| Neither happy<br>nor unhappy | 5 | 3 | 6 | 11 | 14 | 12 |
| Slightly<br>unhappy | 3 | 7 | 6 | 2 | 11 | 4 |
| Moderately<br>unhappy | 6 | 4 | 2 | 6 | 8 | 7 |
| Extremely<br>unhappy | 1 | 2 | 2 | 1 | 10 | 5 |
| Blank | 3 | 3 | 4 | 3 | 3 | 3 |

On the negative side, 18% of students do not feel that they are valued members of the medical school community – They report being moderately or extremely unhappy in this regard. In addition, 12% of students feel similarly dissatisfied in being part of the university community.

Final comments concerning student satisfaction centered around stipend levels and the potential for positive impact of pandemic related disruptions. It was suggested that programs apply lessons learned from the pandemic rather than simply returning to prior, potentially unhealthy, ways of doing things. Finding ways to rectify mismatches in mentor-mentee relationships was noted. Other comments highlighted difficulties due to academic load, lack of communication between departments and students (especially those new to a department), and witnessing unethical behaviors by faculty members. Some students recommended anonymizing this survey to increase participation, while others perceived the university’s response to the Black Lives Matter movement as inadequate. Still others believe graduate student unionization will assist in creating communal feelings. Also mentioned was the need for more social events organized by programs and the school to recognize rising 2^nd^ year students, who have spent their time in graduate school during the worst of the pandemic.

## Conclusion

This survey examined many features of the learning environment that impact on career preparation in the biomedical sciences. Most students are focused on preparing to engage in their future careers. They worry that the pandemic has slowed productivity, limited the ability to grow their professional networks, and produced negative consequences for future employment opportunities. Student career goals and interests remain broad, making clear the need to continue growing career development opportunities.

Students report generally positive relationships with their mentors, although the pandemic has highlighted some concerns regarding mentor expectations and lack of empathy. Overall, a majority of students have positive opinions of their advisors, programs, and departments, while feeling less connected to the medical school, and the university at large.

As the pandemic persists, student comments reveal that its impact on wellness continues to intensify. Although some individuals demonstrate amazing resilience by finding ways to create positive outlets, others are clearly suffering from the pain arising from social isolation and uncertainty in their academic and personal lives. This feedback is critical for guiding further efforts to provide resources, support, and effective training, with the goal of producing positive climate change.

## Acknowledgments

This work was supported by grant CGT025 from the Burroughs Wellcome Fund and by institutional support from the University of Pittsburgh School of Medicine.

## Contributions and Competing Interests

Both authors contributed to all phases of the survey and to writing this report. Neither author has any competing interests to declare.

